# Predicting life-history traits in a stored bean petst beetle *Callosobruchus chinensis* (Coleoptera: Chrysomelidae: Bruchinae) using machine learning

**DOI:** 10.64898/2026.03.07.710260

**Authors:** Xiangpeng Gu, Midori Tuda

## Abstract

Life-history traits play an important role in insect population dynamics and ecological processes. The azuki bean beetle *Callosobruchus chinensis* is a common pest of stored legumes and is also widely used as a model species in ecological and evolutionary research. In this study, we tested whether machine learning models could be used to estimate several traits of *C. chinensis*, including elytral length, development time and adult lifespan.

Experimental data were obtained from laboratory populations. The dataset included biological and environmental variables such as strain, treatment condition, developmental day, sex, temperature, and CO_2_. Six different machine learning models were tested, including linear regression, random forest, support vector machine (SVM), neural network, gradient boosting and AdaBoost. Model performance was evaluated using cross-validation. The coefficient of determination (R^2^) and root mean square error (RMSE) were used to measure prediction accuracy.

Prediction accuracy differed among traits. Elytral length showed relatively higher predictability than the other traits, while development time was difficult to estimate in most models. Lifespan was easier to predict than the other traits, and the neural network produced one of the highest prediction accuracies among the tested models. Feature importance analysis also showed that factors such as sex and treatment condition contributed to variation in several traits.

Machine learning models therefore helped reveal relationships among biological variables and life-history traits in *C. chinensis*. Combining ecological experiments with machine learning analysis may help improve our understanding of insect traits and may support future studies in insect ecology and pest management.

## 1 Introduction

Life-history traits such as body size, development time and adult lifespan are important in insect populations. These traits affect reproduction, survival and population growth. They can also influence how species interact with each other in ecological communities (Chown and Terblanche, 2006). For example, larger insects often produce more eggs and may compete more successfully for resources. Development time also affects how quickly new generations appear. In some stored-product pests, these traits are influenced by food quality, environmental conditions and population density (Kavallieratos et al., 2020; Bhagarathi and Maharaj, 2023).

The azuki bean beetle *Callosobruchus chinensis* (Coleoptera: Chrysomelidae: Bruchinae) is a common pest of stored legumes. It has also been widely used as a model species in ecological and evolutionary studies. Laboratory populations of this beetle have been used to study population dynamics, host–parasitoid interactions and life-history evolution (Zhang et al., 2021; Zhang et al., 2020; Shimada and Tuda, 1996; Vuts et al., 2024). Earlier work showed that development schedules and density effects can lead to oscillating population patterns in experimental populations of *C. chinensis* (Shimada and Tuda, 1996). Other studies found that traits such as development time, adult body size and reproductive output can change with environmental conditions, especially temperature and host bean quality (Tuda and Shimada, 2005). Body size differences may also affect energy storage and reproductive allocation in *C. chinensis* (Yanagi and Tuda, 2012).

In recent years, machine learning methods have been used more often in biological research. These methods can help analyse datasets that contain many variables. Many machine learning models can describe nonlinear relationships between variables, which are sometimes difficult to capture using traditional statistical models. Because of this, machine learning has been used in many ecological and agricultural studies. One common use is insect identification from images. For example, convolutional neural networks have been used to identify insect species from photographs with very high accuracy (Valan et al., 2019). Xia et al. (2018) also developed a neural network that can detect and classify insect species using image data. Machine learning has also been used in pest monitoring systems. For example, automated systems have been developed to detect moth species from trap images to support pest management (Ding and Taylor, 2016). In stored-product pest research, hyperspectral imaging combined with machine learning has been used to distinguish closely related weevil species such as *Sitophilus oryzae* and *Sitophilus zeamais* (Cao et al., 2015).

Machine learning has also been applied to classify insect species and sex. Tuda and Luna-Maldonado (2020) developed supervised learning models using image datasets of the *C. chinensis* together with two parasitoid wasp species. Their study compared several algorithms, including logistic model trees, random forest, support vector machines, multilayer perceptron neural networks and AdaBoost. The models achieved classification accuracies between 88.5% and 98.5%, depending on the task.

However, most studies using machine learning in entomology focus on classification tasks such as species identification or sex recognition. Fewer studies have tried to predict quantitative life-history traits. Predicting traits such as body size, development time and lifespan may help improve our understanding of insect ecology and population dynamics.

In this study, we applied several machine learning models to predict life-history traits of the *C. chinensis*. Six models were tested: linear regression, random forest, support vector machine, neural network, gradient boosting and AdaBoost. These models were used to predict three traits: elytral length, development time and adult lifespan. Model performance was compared using statistical metrics, and the importance of predictor variables was also examined.

## 2 Materials and Methods

### 2.1 Study species

*C. chinensis* is a common pest of stored legumes. It is also widely used in ecological and evolutionary studies. This beetle often attacks seeds of legumes such as azuki bean (*Vigna angularis*), mung bean (*Vigna radiata*), and other stored pulses. Because the life cycle is short and the insects reproduce quickly, the species is easy to maintain in laboratory cultures. For these reasons, *C. chinensis* has been used in many studies on life-history traits, population dynamics and host–parasitoid interactions (Tuda and Shimada, 2005). Females lay eggs on the surface of host beans. The larvae then develop inside the bean and remain there until adult emergence. Traits such as body size, development time and adult lifespan can change depending on environmental conditions, for example, temperature, host quality or population density (Vrtílek et al., 2021). These characteristics make *C. chinensis* a useful system for studying variation in insect life-history traits.

### 2.2 Experimental design and data collection

Newly emerged adults of *C. chinensis* were used in the experiment. Only individuals that had emerged within 3 hours were selected. Each individual was placed in a 5-cm Petri dish containing 5 g of azuki beans, which served as the egg-laying substrate. The Petri dishes were kept in incubators under three conditions (30 °C with 420 ppm CO_2_; 32 °C with 420 ppm CO_2_; 30 °C with 1200 ppm CO_2_). Each treatment had 5 replicates, and relative humidity was kept at about 60–70%.

Eggs were collected during the first 3 days after adult emergence (Day 1; Day 2; Day 3). On each day, ten eggs were randomly selected. To avoid competition among larvae within the same bean, only beans that carried a single egg were used. Each bean with one egg was placed into a separate well of a microplate and labeled. The microplates were then returned to the incubators under the same conditions and kept there until the adults emerged and died. For each individual, several variables were recorded (strain; egg length; development time; sex; elytral length; adult lifespan). These measurements were later used to build the dataset for statistical analysis and machine learning models.

### 2.3 Trait correlation analysis

Relationships among traits were examined using the measured variables. Pearson correlation coefficients were calculated, and the results were displayed as a heatmap. This step allowed us to see whether traits such as body size, development time and lifespan tended to change together.

### 2.4 Machine learning models

Machine learning models were used to predict life-history traits of *C. chinensis*. Six models were tested in this study: Linear Regression; Random Forest; Support Vector Machine; Neural Network; Gradient Boosting; AdaBoost.

All analyses were carried out in Python using the scikit-learn library (Pedregosa et al., 2011).

#### Linear regression

Linear regression was used as a basic model. It describes a linear relationship between predictors and the response variable. The model can be written as

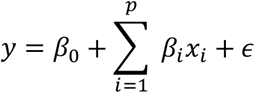

where *y* is the predicted value, *x*_*i*_ are predictor variables, *β*_*i*_ are coefficients and *ϵ* is the error term (James et al., 2023).

#### Random forest

Random forest is a tree-based model that uses many decision trees. Each tree produces a prediction, and the final value is the average of all trees (Breiman, 2001).

The model can be written as

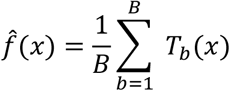

where *T*_*b*_(*x*) is the prediction from the *b*-th tree and *B* is the total number of trees.

#### Support vector machine (SVM)

Support vector machines were also tested. This model separates data using a boundary with the largest possible margin (Hearst et al., 1998). Kernel functions can be used when the relationship between variables is not linear.

#### Neural network

A multilayer perceptron neural network was used as another model. This model contains several layers of connected neurons. The output of a neuron can be written as

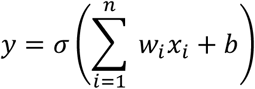

where *w*_*i*_ are weights, *x*_*i*_ are input variables, *b* is the bias term and σ is the activation function (Gurney, 2018).

#### Gradient boosting and AdaBoost

Boosting algorithms improve prediction accuracy by sequentially combining multiple weak learners. AdaBoost iteratively adjusts the weights of training samples to reduce prediction error during model training (Freund and Schapire, 1995). Gradient boosting constructs additive models by fitting new learners to the residual errors of previous models, typically using gradient descent optimization (Friedman, 2001).

### 2.5 Model training

Machine learning models were implemented in Python using the scikit-learn library (Pedregosa et al., 2011). Six algorithms were tested in this study: linear regression, random forest, support vector machine, multilayer perceptron neural network, gradient boosting and AdaBoost. Model performance was evaluated using 10-fold cross-validation. The dataset was randomly divided into ten subsets. In each iteration, nine subsets were used to train the model and the remaining subset was used for validation. This process was repeated ten times so that each subset was used once for validation. For each trait, models were trained using all remaining variables as predictors. The coefficient of determination (R^2^) obtained from cross-validation was used to evaluate predictive performance. Mean R^2^ values and standard deviations were calculated across the ten folds for each model.

### 2.6 Model evaluation

Model performance was evaluated using the coefficient of determination (R^2^) and root mean square error (RMSE).

#### Coefficient of determination

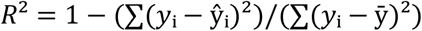

where *y*_i_ is the observed value, ŷ_i_ is the predicted value and 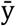 is the mean of the observed values.

#### Root mean square error

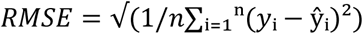

These metrics were used to compare model performance among models and traits. The R^2^ values obtained from each fold were used for the ANOVA test. Differences among models were further examined using one-way ANOVA followed by Tukey’s HSD test. The prediction performance based on R^2^ and RMSE is shown in Figure 2.

### 2.7 Feature importance analysis

Feature importance was calculated using the random forest model to examine which variables affected the predictions most strongly.

This value shows how much each variable contributes to the model. The importance score is based on how much prediction error decreases when a variable is used to split the data in the trees. The results of this analysis are shown in Figure 5.

## 3 Results

### 3.1 Descriptive statistics of life-history traits

A total of 838 individuals of *C. chinensis* were included in the analysis. Several biological variables were recorded for each insect, and three traits were selected as target variables for the prediction analyses: elytral length, development time and adult lifespan. The beetles were reared under different temperature and CO_2_ conditions.

Basic descriptive statistics for these traits are presented in Table 1.

**Table 1.** Summary statistics of three life-history traits in *C. chinensis*.

|  | Elytral length (mm) | Development time (days) | Lifespan (days) |
| --- | --- | --- | --- |
| n | 838 | 838 | 838 |
| mean | 1.879 | 22.104 | 9.612 |
| SD | 0.074 | 1.345 | 1.978 |
| min | 1.572 | 20 | 3 |
| 25% | 1.822 | 21 | 8 |
| 50% | 1.872 | 22 | 9 |
| 75% | 1.942 | 23 | 11 |
| max | 2.042 | 29 | 18 |

Note: The table shows descriptive statistics for three life-history traits measured in *C. chinensis*. Elytral length represents adult body size and is measured in millimeters (mm). Development time refers to the number of days from egg to adult emergence. Lifespan represents adult longevity and is measured as the number of days an individual survived after emergence. Values include the sample size (n), mean, standard deviation (SD), minimum value (min), quartiles (25%, 50%, 75%), and maximum value (max).

Adult lifespan varied considerably among individuals in the dataset. Development time also differed among individuals, although the variation was somewhat smaller than that observed for lifespan.

Adult body size was represented by elytral length. This trait also showed variation within the population, although the overall range was moderate. Variation in body size is commonly observed in *C. chinensis* and may be associated with several ecological factors such as larval competition within host beans, differences in bean quality and environmental conditions during development (Yanagi and Tuda, 2012; Martinossi-Allibert et al., 2019).

Previous studies have also reported that body size can be related to other life-history traits in *C. chinensis*. For example, body size may influence reproductive allocation and energy storage during development (Yanagi and Tuda, 2012). Environmental stress during development can also affect traits such as development time and survival (Maharjan et al., 2017; Sun et al., 2024).

Overall, the dataset shows clear variation among individuals for the traits examined. This variation provides useful information for building predictive models of life-history traits.

### 3.2 Trait correlations

Relationships among the measured traits (elytral length; development time; adult lifespan) were examined using Pearson correlation analysis. The results are shown as a correlation heatmap in Figure 1.

**Figure 1.**
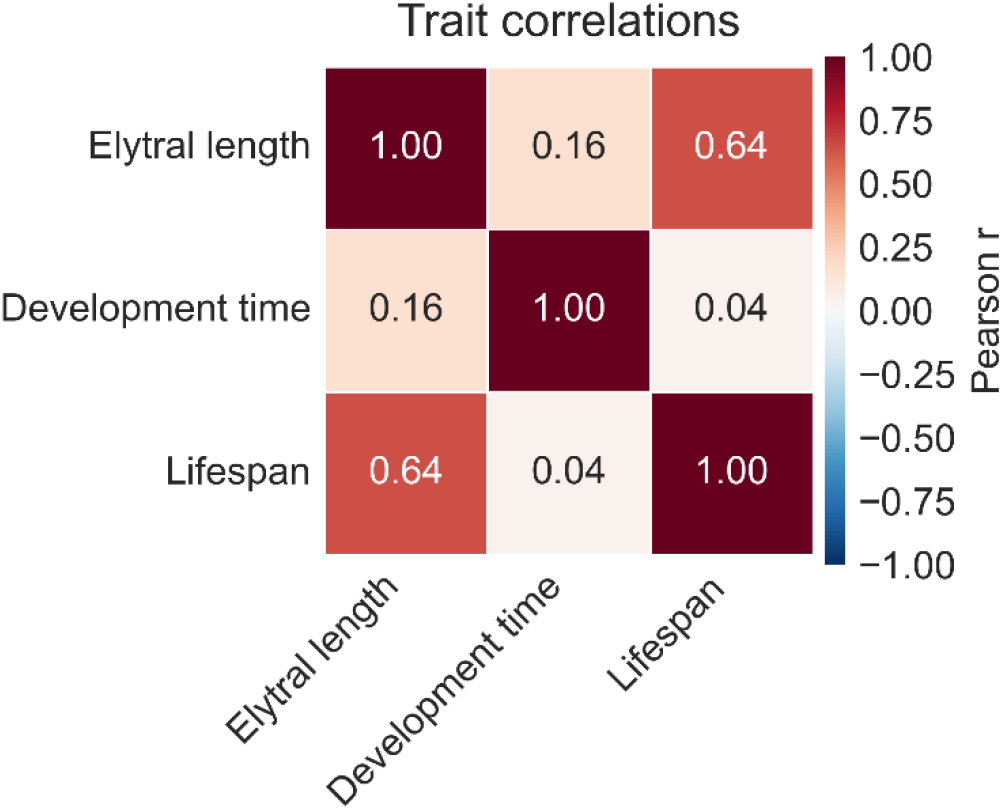
Correlations among life-history traits of *C. chinensis*. Pearson correlation matrix showing the relationships among three life-history traits: elytral length, development time and adult lifespan. Values represent Pearson correlation coefficients (r). Warmer colors indicate stronger positive correlations, while cooler colors represent weaker correlations.

Some patterns were observed among the traits. Elytral length showed a moderate positive correlation with adult lifespan (r = 0.64). Individuals with larger body size tended to have longer adult lifespan.

In contrast, development time showed only weak relationships with the other traits. The correlation between development time and elytral length was low (r = 0.16), and the association between development time and lifespan was very weak (r = 0.04).

These results indicate that body size and adult lifespan may be partially related, whereas development time appears to vary more independently from the other traits.

### 3.3 Comparison of machine learning models

Six models were used to predict three traits of *C. chinensis*: elytral length, development time and adult lifespan. The results of the model comparison are shown in Figure 2.

**Figure 2.**
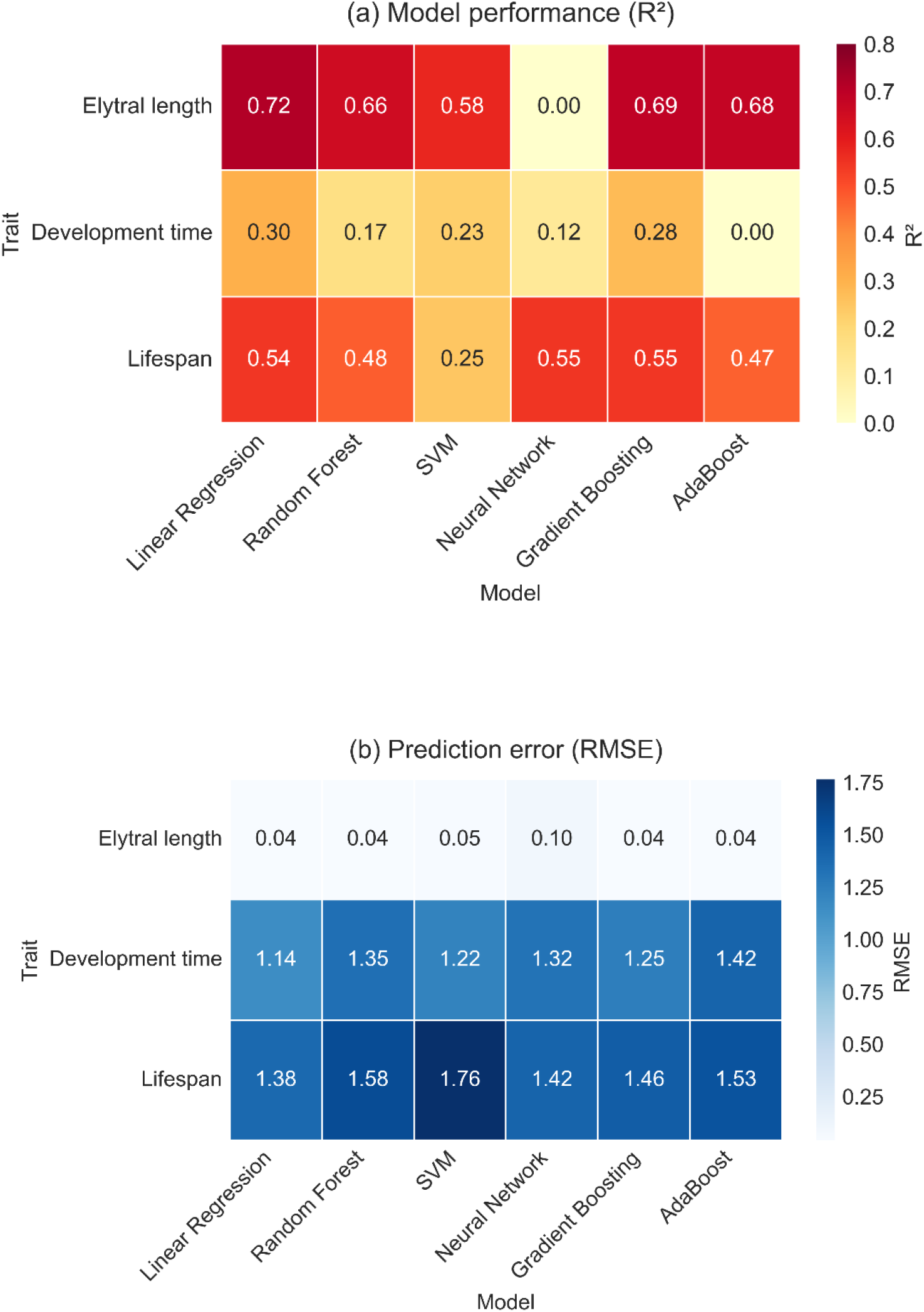
Performance of six machine learning models for predicting life-history traits. Heatmaps showing model performance when predicting elytral length, development time and adult lifespan. (a) Prediction accuracy measured by the coefficient of determination (R^2^). (b) Prediction error measured by the root mean square error (RMSE). Six models were evaluated: linear regression, random forest, support vector machine, neural network, gradient boosting and AdaBoost.

Prediction performance differed among the three traits. Negative R^2^ values obtained during cross-validation were set to 0 for visualization purposes. Elytral length was the easiest trait to predict. Most models produced relatively high R^2^ values for this trait, ranging from about 0.58 to 0.72. Linear regression gave the highest value (R^2^ = 0.72), while gradient boosting and AdaBoost also showed strong performance. This indicates that the models were able to capture variation in elytral length relatively well.

Adult lifespan showed intermediate prediction accuracy. R^2^ values for this trait ranged from about 0.25 to 0.55 across the models. The neural network and gradient boosting models produced the highest values (R^2^ ≈ 0.55). Other models produced slightly lower values but still captured part of the variation in lifespan.

Development time was the most difficult trait to predict. Most models produced relatively low R^2^ values, generally below 0.30. In some cases, the values were close to 0. This suggests that development time may depend on additional factors that were not included among the predictor variables used in the present dataset.

Another observation was that ensemble models did not consistently outperform simpler approaches. For example, linear regression produced the best result for elytral length even though more complex models were also tested. The dataset used in this study was moderate-sized dataset, which may explain why simpler models sometimes performed as well as, or even better than, more complex ones.

Taken together, prediction success differed among traits. Elytral length showed the highest predictability, lifespan showed moderate predictability, and development time was the most difficult trait to estimate.

Mean R^2^ values of the six models are shown in Figure 3. Error bars represent standard deviation, and different letters indicate significant differences among models based on Tukey’s HSD test.

**Figure 3.**
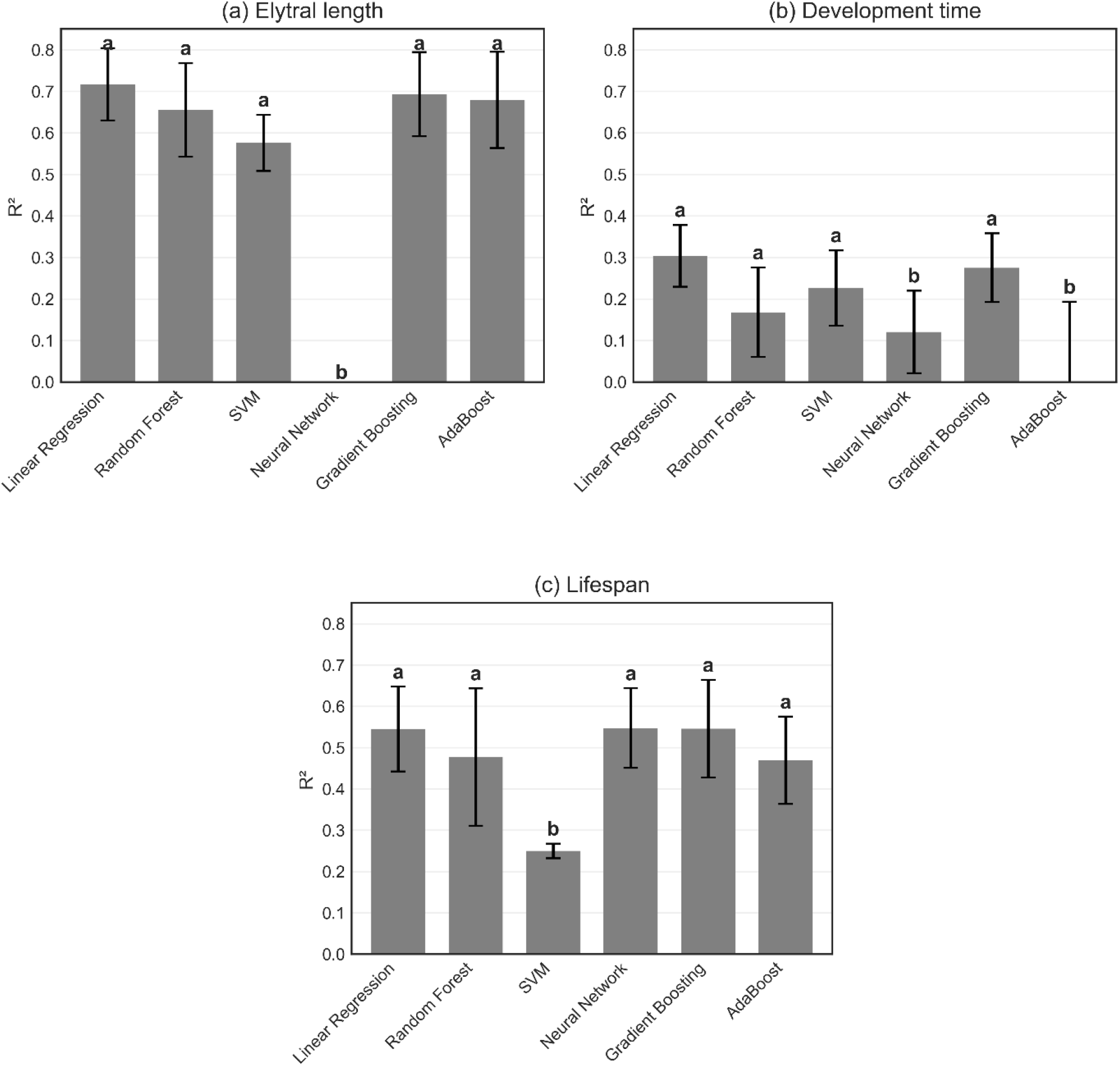
Comparison of prediction accuracy among machine learning models. Mean R^2^ values obtained from cross-validation for each model when predicting (a) elytral length, (b) development time and (c) adult lifespan. Error bars represent standard deviations. Different letters above bars indicate statistically significant differences among models.

### 3.4 Prediction performance of machine learning models

Predicted values were compared with the observed values for the three traits. The relationships between predicted and observed values are shown in Figure 4.

**Figure 4.**
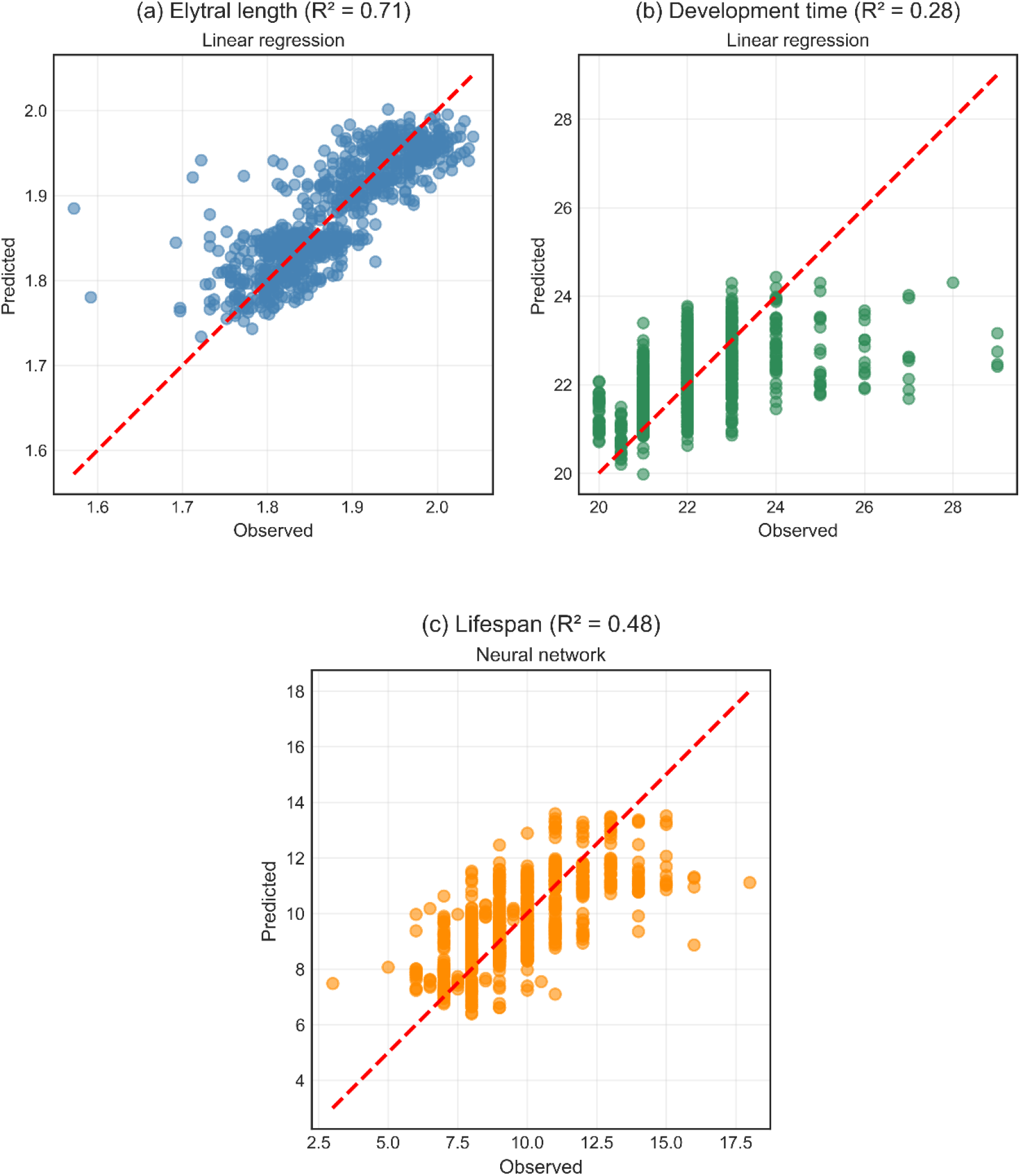
Observed and predicted values for the best-performing models. Scatter plots comparing observed and predicted values for the three traits. (a) Elytral length predicted using linear regression. (b) Development time predicted using linear regression. (c) Adult lifespan predicted using the neural network model. The dashed red line indicates the 1:1 relationship between observed and predicted values.

For elytral length, most predicted values were close to the 1:1 reference line. The points are concentrated near the line, and predicted values are generally similar to the observed ones. The relatively high R^2^ value reported in the model comparison (Figure 2) shows the same pattern.

For adult lifespan, predicted values also increase as observed values increase. The points are more scattered around the reference line than for elytral length, which reflects lower prediction accuracy.

For development time, the relationship between predicted and observed values is weaker. A positive trend can still be seen, but the points are spread more widely around the line than for the other two traits.

The three panels therefore show different levels of agreement between predicted and observed values. Elytral length shows the closest agreement, lifespan shows moderate agreement, and development time shows the largest spread of points.

### 3.5 Feature importance analysis

Feature importance was calculated using the random forest model to examine which variables contributed most to the predictions. The relative importance of each predictor is shown in Figure 5.

**Figure 5.**
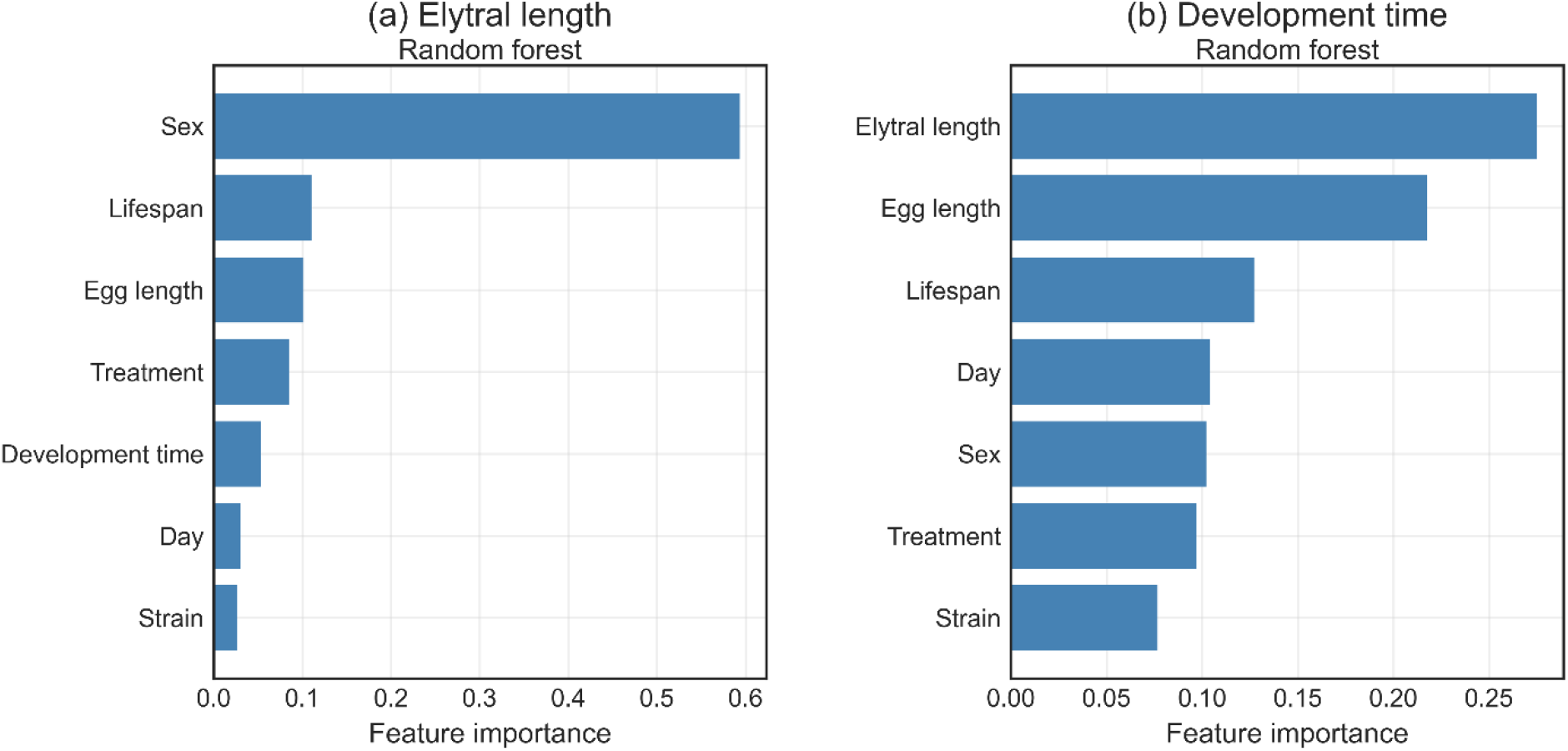
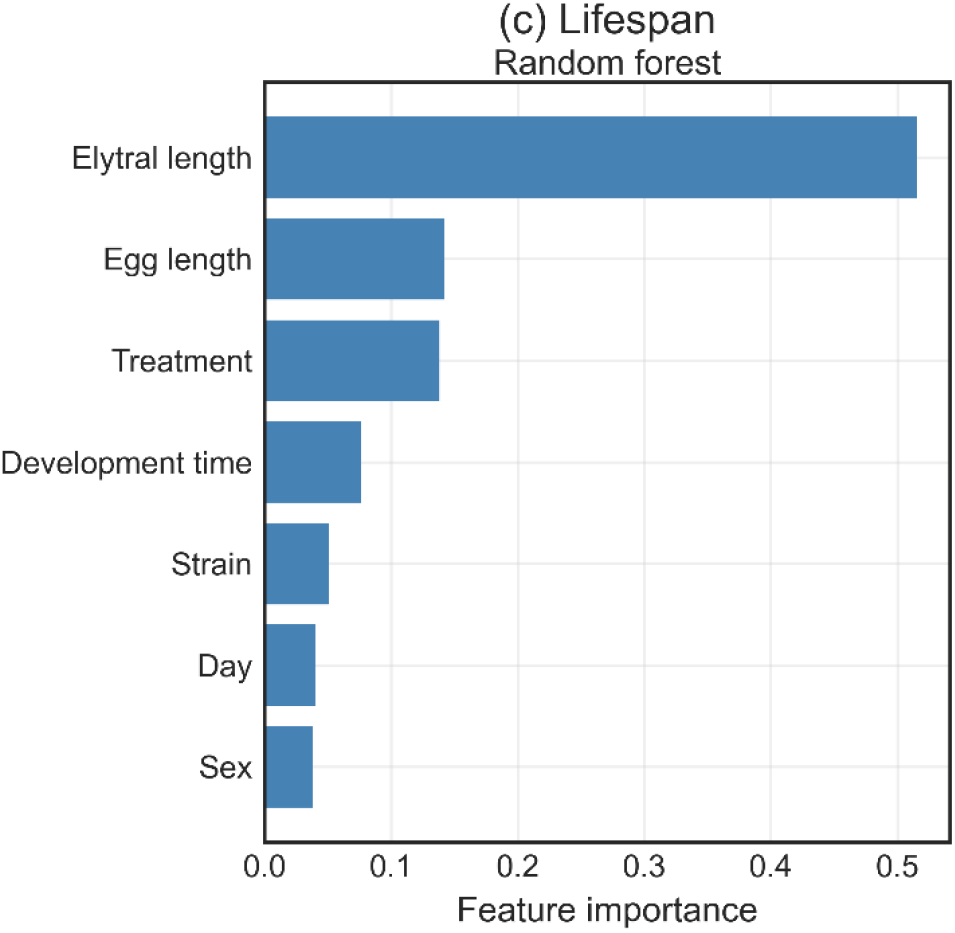
Relative importance of predictor variables in random forest models. Feature importance values indicating the contribution of each predictor variable to model predictions. Separate models were fitted for (a) elytral length, (b) development time and (c) adult lifespan. Higher values indicate greater influence of a predictor on model predictions.

The importance patterns were different for the three traits.

For elytral length, sex had by far the highest importance value. In many insects, males and females differ clearly in body size. In *C. chinensis*, females are usually larger than males. Because of this size difference, body size measurements such as elytral length are closely related to sex.

For development time, elytral length and egg length had the highest importance values. Other variables, including lifespan, day, sex and treatment, also contributed to the model, but their values were smaller. Development time in insects can be affected by several factors during growth, such as temperature, food availability and genetic background. Because several factors act together, more than one variable may influence this trait.

For adult lifespan, elytral length had the highest importance value. Egg length and treatment also had moderate importance values, while the remaining variables contributed less. Lifespan in insects is often related to body size and conditions experienced during development.

The three traits therefore showed different importance patterns. Elytral length was mainly linked to sex, while lifespan and development time were influenced by several biological and environmental variables.

### 3.6 Model evaluation and residual analysis

Model performance was also examined using residual plots together with root mean square error (RMSE) values. Residuals were calculated as the difference between observed and predicted values for each trait. Residual plots for the three traits are shown in Figure 6. In most cases, residuals were distributed around zero and no clear directional pattern appeared across the prediction range.

**Figure 6.**
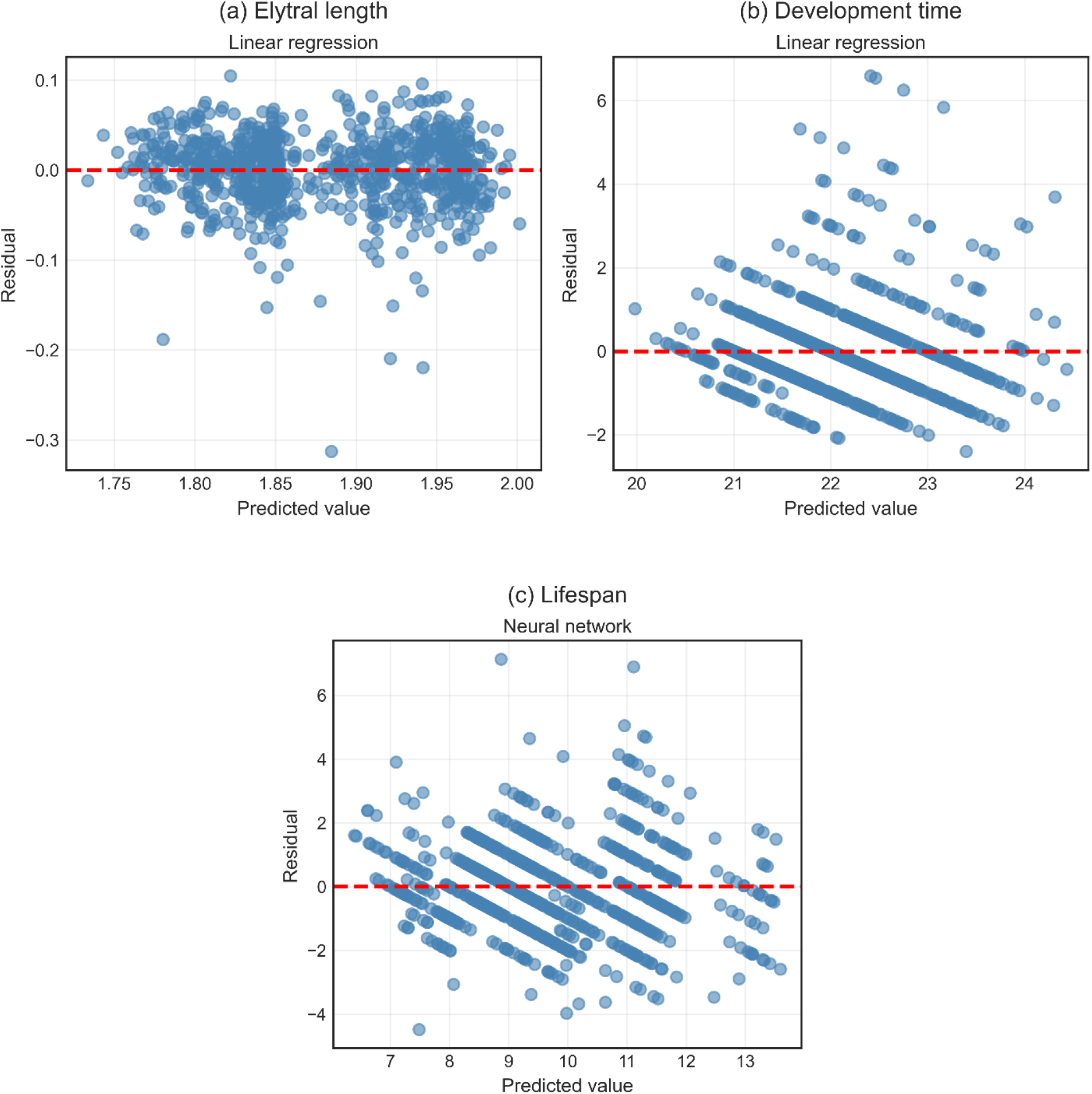
Residual diagnostics of prediction models. Residual plots showing the relationship between predicted values and residual errors for (a) elytral length predicted by linear regression, (b) development time predicted by linear regression, and (c) adult lifespan predicted by the neural network model. The dashed horizontal line indicates 0 residual error.

For elytral length, residual values were generally small and concentrated near 0. This indicates that predicted values were usually close to the observed values. This pattern agrees with the relatively high R^2^ values reported for this trait in Figure 2.

For adult lifespan, residuals showed a wider spread than for elytral length. Although many points were still located near 0, larger deviations were also present.

For development time, residuals were distributed more broadly across the prediction range. The general pattern was captured by the models, but a larger amount of variation remained unexplained.

The RMSE values showed a similar tendency (Figure 2). Prediction errors were smallest for elytral length, followed by adult lifespan, while development time showed the largest prediction errors.

## 4 Discussion

### 4.1 Predictability of life-history traits in *C. chinensis*

In this study we examined whether several life-history traits of *C. chinensis* could be predicted using machine learning models. The three target traits were elytral length, development time and adult lifespan. The results showed clear differences in how well these traits could be predicted.

Among the three traits, elytral length showed the highest prediction accuracy. In several models the R^2^ values were relatively high, reaching about 0.72 in the best case. Adult lifespan showed moderate prediction accuracy, with the highest R^2^ value around 0.55. Development time was much harder to predict, and most models produced lower R^2^ values, usually below 0.30. These differences suggest that some biological traits can be estimated more easily from the available variables than others.

One reason why elytral length was predicted relatively well may be the strong effect of sex on body size in *C. chinensis*. In many insect species males and females differ clearly in body size. Females often need larger body size because egg production requires space for reproductive tissues and stored resources (Cueva del Castillo, 2015). In *C. chinensis*, females are generally larger than males (Singh et al., 2024; Premkumari et al., 2023). As a result, body size traits such as elytral length often show a consistent difference between the sexes (Benítez et al., 2020; Ferracini et al., 2025). The feature importance analysis in this study also showed that sex contributed most strongly to the prediction of elytral length. Because this relationship is quite stable, the models were able to capture a large part of the variation in this trait.

Adult lifespan showed a moderate level of predictability. The neural network produced one of the highest R^2^ values, although several other models gave similar results. Lifespan in insects is often influenced by the resources obtained during larval development. Individuals that receive more nutrition during development may store more energy and therefore survive longer after emergence (Yan et al., 2021; Zhang et al., 2025; Fernández et al., 2025). These processes can create links between lifespan and other biological traits measured in the dataset.

*C. chinensis* illustrates this relationship well. Their larvae develop entirely inside host beans, and the amount of food available within each bean can influence adult traits such as lifespan and fecundity. Earlier studies have shown that host quality and resource availability during larval development can affect adult survival and reproductive output in *C. chinensis* (Mounika et al., 1997; Yang et al., 2025). Because some of these biological processes are indirectly reflected in variables such as body size, the models used in this study were able to capture part of the variation in adult lifespan.

Development time showed the lowest prediction accuracy among the three traits. Developmental duration in insects is affected by several interacting factors, including temperature, resource availability, genetic background and other developmental conditions (Vasudeva, 2023; Martinossi-Allibert et al., 2018; Mackay et al., 2023). Some of these factors were not directly included in the present dataset. For this reason, the models were only able to explain a limited proportion of the variation in development time.

Taken together, the results suggest that morphological traits such as body size may be easier to estimate from biological variables, while developmental traits influenced by many environmental factors may be more difficult to predict.

### 4.2 Prediction of morphological traits

Among the three traits examined in this study, elytral length showed the highest prediction accuracy. Most of the tested models produced relatively high R^2^ values (Figure 2), which indicates that body size variation in the dataset could be captured reasonably well by the predictor variables.

A likely explanation is the strong influence of sex on body size in *C. chinensis*. Sexual size dimorphism is common in many insect species. Females often require larger body size to support egg production and reproductive functions, while males may benefit from smaller body size that allows greater mobility (Zhu et al., 2025). In *C. chinensis*, females are typically larger than males. This difference produces clear variation in body size measurements such as elytral length.

The feature importance analysis supports this interpretation. Sex had the highest importance value for predicting elytral length, indicating that this variable contributed strongly to the model predictions (Figure 5). Because the size difference between males and females is relatively consistent, the models can use sex information to estimate body size more accurately.

Body size in insects can also be affected by environmental conditions during development. Factors such as host bean quality, larval competition and temperature may influence growth and final adult size. For example, in the mosquito *Anopheles gambiae*, higher temperatures during larval and adult stages were shown to reduce adult longevity and alter reproductive traits such as egg production and mating success (Christiansen-Jucht et al., 2015). In addition, experiments on anuran larvae showed that higher temperatures and better food quality increased larval growth rate and accelerated metamorphosis, while low food quality slowed development and resulted in smaller body size (Álvarez and Nicieza, 2002). Therefore, when several larvae develop within the same bean, competition for resources can also reduce the amount of food available to each individual and lead to smaller adult size.

Temperature may further influence body size through its effect on development rate. In many ectothermic organisms, higher temperatures accelerate development but shorten the growth period. As a result, individuals that develop at higher temperatures often become smaller adults. This pattern is commonly known as the temperature–size rule (Angilletta et al., 2003; Forster and Hirst, 2012).

Although these factors can produce variation in body size, morphological traits such as elytral length may still be relatively predictable when strong biological patterns are present (Barton, 2011). In the present dataset, the clear size difference between males and females likely contributed to the relatively high prediction accuracy observed for this trait.

### 4.3 Biological relationships among traits

Life-history traits are often interconnected because they are influenced by shared physiological and developmental processes (Braendle et al., 2011; Leyria et al., 2025). The results of this study provide several examples of such relationships.

The correlation analysis showed a moderate positive relationship between elytral length and adult lifespan (r = 0.64) (Figure 1). Individuals with larger body size tended to live longer. This relationship is consistent with the feature importance analysis, which showed that elytral length was one of the most important predictors of adult lifespan.

One possible explanation is that larger individuals often possess greater energy reserves. During larval development, insects accumulate resources that are later used to support adult survival and reproduction. Larger body size may therefore reflect greater energy storage capacity. Individuals with larger reserves may survive longer, particularly when environmental conditions become stressful (Martinossi-Allibert et al., 2019).

Similar relationships between body size and life-history traits have been reported in previous studies on *C. chinensis*. Yanagi and Tuda (2012) showed that female body size in *C. chinensis* can influence reproductive traits such as egg size. In their study, the size of reproductive organs limited the amount of resources that could be allocated to each egg. As a result, larger females were able to produce larger eggs.

These findings illustrate how morphological traits and reproductive characteristics can be linked through resource allocation processes. Growth, reproduction and survival are therefore not independent traits but are connected through the physiological constraints experienced by organisms during development.

Because of these complex relationships, analytical approaches that can consider multiple variables simultaneously may be particularly useful for studying life-history traits. Machine learning models provide one such approach, as they can incorporate multiple predictors and capture nonlinear relationships among biological variables.

### 4.4 Applications of machine learning in insect ecology

Machine learning methods are increasingly used in ecological research. In many studies these approaches are applied to tasks such as insect identification, pest monitoring and analysis of large image datasets (Wang et al., 2025; Høye et al., 2021).

For example, image-based classification has been widely used to identify insect species and determine sex. Recent studies using deep learning models have shown that insect species and gender can be identified with very high accuracy from image data. For instance, a deep learning system based on the YOLO algorithm successfully classified mosquito species and sex with high performance, achieving a mean average precision of 99% and sensitivity of 92.4% (Kittichai et al., 2021). These results demonstrate that machine learning approaches can provide powerful tools for insect identification and monitoring. These studies illustrate how artificial intelligence techniques can help analyse biological data. Machine learning models are particularly useful when relationships among variables are complex or nonlinear and may not be easily captured by traditional statistical approaches.

In the present study, a similar approach was applied to the prediction of life-history traits rather than classification tasks. The results show that machine learning models can also help explore relationships among biological traits in insect populations. Although prediction accuracy differed among traits, the models were still able to detect meaningful biological patterns in the dataset.

These findings suggest that machine learning can serve as a complementary tool for ecological research. When combined with experimental data, such methods may help reveal patterns that are difficult to identify using conventional analytical approaches.

### 4.5 Limitations and future directions

Several limitations of the present study should be considered.

First, the number of predictor variables in the dataset was relatively limited. Additional environmental information, such as bean size, bean nutritional content or more detailed micro-environmental conditions, could potentially improve prediction accuracy. These variables may influence larval growth and adult traits but were not directly measured in the current experiment.

Second, dataset size may also affect model performance. Machine learning models generally perform better when larger datasets are available, because larger datasets allow algorithms to identify patterns more reliably and reduce the risk of overfitting (Albattah and Khan, 2025; Naser, 2026).

Third, future studies could examine additional modelling approaches. More advanced methods, including deep learning models or alternative ensemble algorithms, may further improve predictive performance when larger datasets are available.

Despite these limitations, the present study shows that combining ecological experiments with machine learning analysis can provide useful insights into insect life-history traits. Integrating biological datasets with modern computational methods may therefore open new opportunities for studying insect population dynamics and life-history strategies.

## 5 Conclusion

In this study we examined whether machine learning models could be used to predict several traits of *C. chinensis*. The dataset included measurements of egg length, elytral length, development time and adult lifespan collected under different environmental conditions. Six machine learning models were tested and compared.

Prediction accuracy differed among the three target traits. Elytral length showed the highest prediction accuracy, with the best model reaching an R^2^ value of about 0.72. Adult lifespan showed moderate prediction performance, with the highest R^2^ value around 0.55. Development time was the most difficult trait to predict and generally produced lower R^2^ values.

The analysis of variable importance also revealed several biological patterns. Sex had a strong influence on elytral length, which is consistent with the clear body size difference between males and females in *C. chinensis*. Elytral length and several other biological traits also contributed to the prediction of adult lifespan.

These results suggest that machine learning models can help explore relationships among biological traits in insect populations. Combining ecological experiments with machine learning analysis may improve our understanding of insect life-history patterns and may also support future studies on integrated pest management (IPM).

